# Microencapsulation of a novel *Bacteroides thetaiotaomicron* strain: a promising strategy to fortify intestinal barrier fortification in weaned pig model

**DOI:** 10.64898/2026.03.11.711050

**Authors:** Shen Jin, Yanjiao Liu, Yifang Zhang, Yuqing Shen, Cong Lan, Hua Li, Jun He, Aimin Wu, Jiayong Tang, Ruinan Zhang, Huifen Wang, Quyuan Wang, Gang Tian, Jingyi Cai, Xiangbing Mao, Edward Liam Good, Yuheng Luo

## Abstract

Porcine *Bacteroides thetaiotaomicron* LYH5 demonstrated in vitro antimicrobial activity, suggesting probiotic potential. Due to poor gastric juice tolerance, LYH5 was encapsulated via extrusion using sodium alginate (SA) and gellan gum. Box-Behnken design optimization yielded optimal parameters: SA 1.5%, gellan gum 0.4%, CaCl₂ 0.9%, bacteria:glue ratio 1:4, achieving an encapsulation rate of 84.22±0.17%. Its effect on weaned piglet intestinal health was evaluated using 78 piglets (7.69±0.52 kg) randomly assigned to 4 groups for 40 days: CON (control), T (basal diet + LYH5 live bacteria, 1×10¹⁰ CFU/mL), TJ (basal diet + LYH5 microcapsules, 1×10¹⁰ CFU/mL, J (basal diet + empty capsules). The results of this experiment showed that compared with the control group, LYH5 microcapsule can improve the intestinal barrier function without affecting the growth performance of piglets, and provide ideas and references for the development of human next-generation probiotics (NGP).

**IMPORTANCE:** This study addresses the key bottleneck of poor gastric acid tolerance of probiotics via microencapsulation and provides a practical reference for the development of human next-generation probiotics.

## INTRODUCTION

*Bacteroides* spp. represent the most abundant gram-negative bacteria in the human intestinal tract, with concentrations reaching 10^9^-10^11^ CFU per gram of feces (60). Notabley, this genus is a core component of the monogastric animal gut microbiota, particularly enriched in the colonic mucus layer (17). In piglets, *Bacteroides* dominate the fecal microbiome before and after weaning, comprising over 90% of the bacterial community (30). According to our previous research, reduced abundance of *Bacteroides* has been observed in diarrheic weaned piglets, suggesting its potential as a microbial marker for diarrhea and its association with impaired carbohydrate/polysaccharide metabolism (44).

Among the *Bacteroides* species, *Bacteroides thetaiotaomicron* is particularly noteworthy. This species constitutes 6% of total fecal bacteria in healthy adults and 12% of the *Bacteroides* community (54). It has been widely used in genetic modification research and serves as a model organism for studying human health and diseases (61). Like other *Bacteroides*, *B. thetaiotaomicron* metabolizes polysaccharides to produce short-chain fatty acids (SCFAs), which are absorbed by the host (43). As a paradigm of polysaccharide utilization, it can exploit both dietary and host-derived polysaccharides, a trait that has enabled its partial application in pharmaceutical and food industries (57). Its genome-encoded proteins and secreted enzymes facilitate sugar capture in the colon, providing substrates for membrane-bound glycoside hydrolases to effectively utilize carbohydrates (25). Recent studies have shown that *B. thetaiotaomicron* DSM 2079 promotes the growth of butyric acid-producing bacteria via lactose metabolism (10), underscoring its role in juvenile intestinal health. While its effect on intestinal inflammation, particularly colitis, has been explored in preclinical models, such as reducing colonic lesions in IL-10 knockout mice (15), mechanistic studies remain limited. Beyond the bacteria themselves, *B. thetaiotaomicron* metabolites plays a critical role in host gut health. For instance, acetic acid produced by *B. thetaiotaomicron* VPI-5482 indirectly supports the growth of *Faecalibacterium prausnitzii* by serving as a substrate, highlighting inter-bacterial ecological balance and its importance for colonic epithelial homeostasis (53). Additionally, *B. thetaiotaomicron*-derived indoleacetic acid (IAA) and indole propionic acid (IPA), agonists of the aromatics receptor (AHR), alleviate colonic inflammation by activating AHR and regulating CD4^+^ T cell differentiation (34). In LPS-challenged mice, its motabolic oleic acid reduces the expression of pro-inflammatory markers such as COX-2, TNF-α, IL-6 and IL-12 in macrophages (6). Most NGPs are highly sensitive to oxygen, gastric acid and bile salts, limiting their therapeutic application (56). Microencapsulation technology offers a solution, though its use in NGPs remains in its infancy. Furthermore, the functions of most probiotics exhibit significant strain-specificity (28) and host-specificity (3, 37), while research on NGPs suitable for weaned piglets remains extremely limited.

Given this, our study first isolated a probiotic potential *Bacteroides* strain from fresh feces of healthy weaned piglets. We then enhanced its tolerance to gastric juice and intestinal digestive fluids via microencapsulation technology, and preliminarily evaluated its promotion effect on intestinal health in weaned piglets. Our work may provide novel insights into microbiota screening for human intestinal health research.

## MATERIALS AND METHODS

### Bacterial strains

*B. thetaiotaomicron* strain used in this study was isolated from fresh feces of healthy weaned piglets and deposited in the China Center for Type Culture Collection (CCTCC) under the accession number CCTCC NO: M2021752. *Enterotoxigenic Escherichia coli* ATCC 25922, *Salmonella cholerae suis* ATCC 527 and *Staphylococcus aureus* ATCC 25923 were maintained in our laboratory culture collection.

### Identification of porcine *B. thetaiotaomicron* strain

Genomic DNA of *B. thetaiotaomicron* LYH5 was extracted using the protocol previously established by our research group (38). The bacterial 16S rDNA was amplified by PCR using universal primers 27F (5’-AGAGTTTGATCMTGGCTCAG-3’) and 1492R (5’-TACGGYTACCTTGTTACGACTT-3’). The PCR program consisted of initial denaturation at 94 ℃ for 5 min, 30 cycles of denaturation at 94 ℃ for 30 s), annealing at 50 ℃ for 30 s, extension at 72 ℃ for 30 s, and a final extension at 72 ℃ for 5 min. The amplified product was subjected to Sanger sequencing by Sangon Biotech (Shanghai) Co., Ltd. (China), and the obtained gene sequence was aligned with 16S rRNA sequences in NCBI GenBank (https://blast.ncbi.nlm.nih.gov/) to identify the most similar strain.

For morphological characterization, the isolated strain was streaked onto modified solid Gifu Anaerobic Medium (GAM) and anaerobically cultured at 37°C for 48 h, followed by scanning electron microscopy analysis performed by Chengdu Lilai Biotechnology Co., Ltd. Physiological and biochemical properties were evaluated via Gram staining, catalase test, gelatin liquefaction test, and motility test. Hemolytic activity was assessed by streaking the strain on sheep blood agar plates and observing hemolysis after 48 h of anaerobic incubation. Antibiotic resistance was determined using Oxoid antibiotic discs. Sixteen antibiotics were placed on modified GAM agar plates, and inhibition zones were measured after 48 h of anaerobic culture at 37°C. Resistance was interpreted according to European Food Safety Authority (EFSA) guidelines (1).

### *In vitro* antimicrobial assays

Antimicrobial activity of the isolated strain was evaluated using the agar diffusion Oxford cup assay and metabolite co-culture assay. For the Oxford cup assay, the culture supernatants of the strain were prepared, and test pathogens (*E. coli* ATCC 25922, *S. cholerae suis* ATCC 527, and *S. aureus* ATCC 25923) were adjusted to 10^-6^∼10^-7^CFU/mL in Luria-bertani (LB) broth, then mixed with molten LB agar (45°C) and poured into Petri dishes. After solidification, Oxford cups were placed on the agar surface, and 15 μL of the isolated strain fermentation broth was added to each cup. Plates were incubated at 37°C for 48 h under anerobic conditions, and inhibition zone diameters were measured. For the metabolite co-culture assay, the fermentation broth of the strain was centrifuged at 12,000 × *g* for 10 min at 4°C, and the supernatant was extracted with ethyl acetate (1:1, v/v). The organic phase was collected, dried under nitrogen, resuspended in LB medium, and filtered through a 0.22-μm membrane. The purified metabolites were co-cultured with pathogens, and OD_600_ absorbance was measured every 2 h. Each group was performed in triplicate.

SCFAs including acetic acid, propionic acid, butyric acid, valeric acid, and total SCFAs in culture of the strain were quntified by gas chromatography (GC, Varian CP-3800, flame ionization detector) using a 10 μL manual injector. following our previously established method(38).

Outer membrane vesicles (OMVs) of the isolated strain were isolated by ultracentrifugation. Following 72-h anaerobic culture in modified GAM medium, the bacterial suspension was centrifuged at 150,000 × g for 30 min to remove cells, and the supernatant was filtered through a 0.22-μm membrane. The filtrated was centrifuged again and the pellet was washed twice with PBS, resuspended in PBS, aliquoted, and stored at −80°C. OMVs were visualized by scanning electron microscopy after negative staining with 2% urany acetate.

### Tolerance test of isolated strain

To prepare simulated gastric fluid (SGF), 0.2% NaCl solution was prepared with sterilized deinoized water, adjusted to pH 2.0 using 1 M HCl, and supplemented with 0.32% pepsin (w/v). To prepare simulated intestinal fluid (SIF), 6.8 g/L KH_2_PO_4_ solution was prepared, ajusted to pH 7.4 with NaOH, and supplemented with 10 mg/mL trypsin. Both fluids were stored at 4 ℃, filter-sterilized through 0.22-μm nylon membrane filters prior to use, and preheated to 37℃ before experimentation.

Log-phase culture (1 mL) of the strain were inoculated into sterile anaerobic tubes containing 9 mL of preheated SGF or SIF. The mixtures were incubated at 37 ℃ with shaking (220 rpm), and samples were collected at 0, 10, 20, 30 and 60 min. Bacterial viability was determined by serial dilution plating on modified GAM agar, with triplicate experiments for each time point.

### Preparation of microcapsules

#### Optimization of preparation process parameters

Sodium alginate (SA, Qingdao Mingyue Seaweed Group, China), gellan gum (Xinjiang Fufeng Biotechnology, China), and anhydrous calcium chloride (Chengdu Cologne Chemical Co., Ltd., China) were used. All materials were autoclaved at 121°C for 20 min. Cells of the isolated strain in late logarithmic phase were harvested by centrifugation at 5,000 × *g* for 10 min at 4℃, washed twice with PBS by re-centrifugation, and resuspended in sterile saline to a concentration of 1 × 10^10^ CFU/mL.

Single-factor experiments were first conducted to determine the optimal ranges of SA, gellan gum, calcium chloride, and bacterial-to-gel ratio. The Box-Behnken design of Design-Expert 10 software was used for response surface methodology (RSM) to optimize the microencapsulation conditions. The experimental factor levels (Table 1) comprised 29 test points with 5 central repeats for error estimation.

**Table 1.**
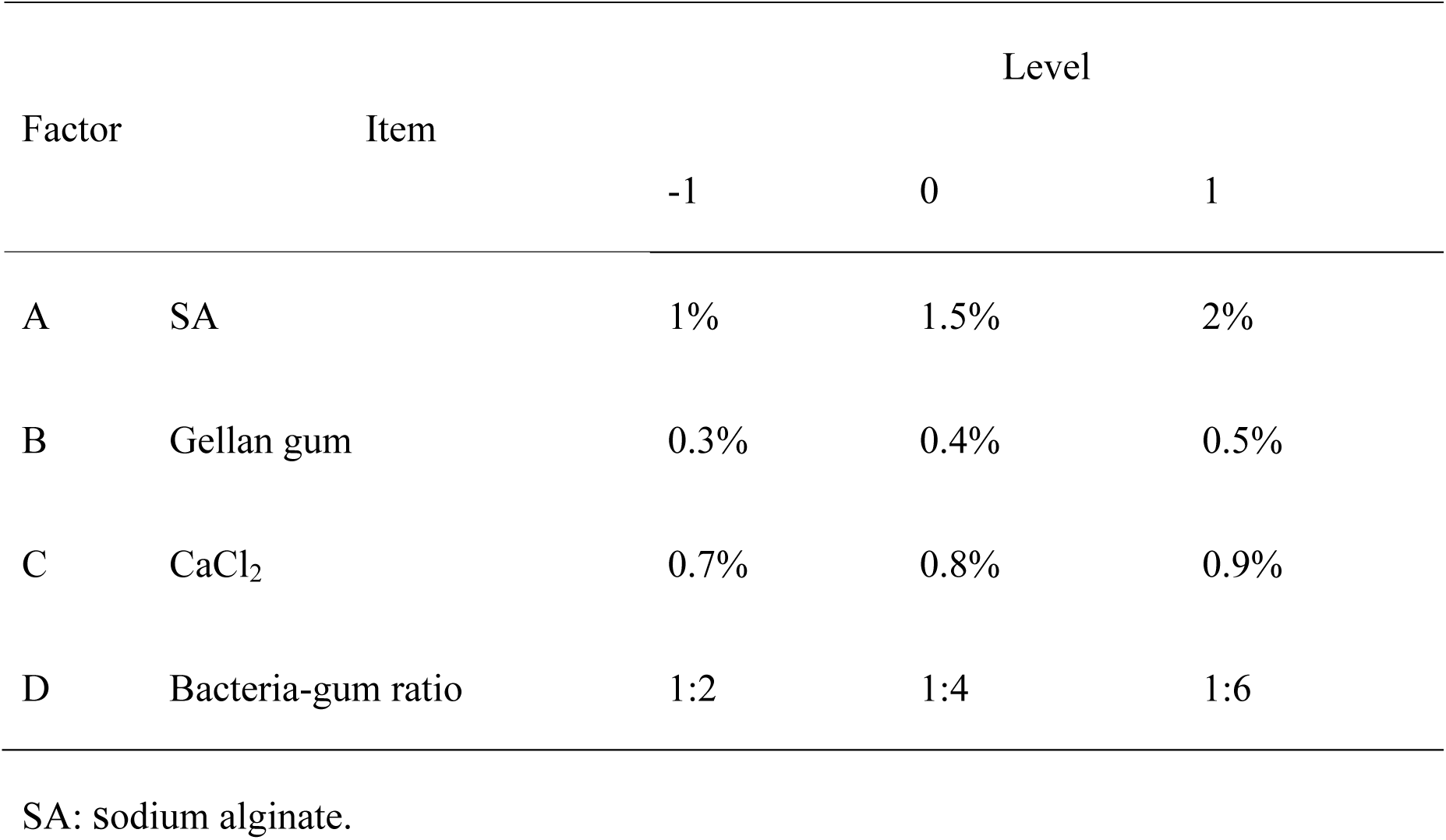
Response factors and levels.

| Factor | Item | Level |  |  |
| --- | --- | --- | --- | --- |
|  |  | -1 | 0 | 1 |
| A | SA | 1% | 1.5% | 2% |
| B | Gellan gum | 0.3% | 0.4% | 0.5% |
| C | CaCl<sub>2</sub> | 0.7% | 0.8% | 0.9% |
| D | Bacteria-gum ratio | 1:2 | 1:4 | 1:6 |

SA: sodium alginate.

Under aseptic and anaerobic conditions, 1 g of wet microcapsules was mixed with 9 mL of pH 7.0 sterile PBS, vortexed, and serially diluted (10-fold) with sterile saline. Aliquots (100 μL) were spread-plated on modified GAM agar, and colonies were counted after 48-h anaerobic incubation at 37℃. The encapsulation efficiency (A, %) was calculated as: A = (a/b) × 100, where a is the viable cell count (CFU) in the microcapsule suspension, and b is the initial viable cell count (CFU) in the bacterial suspension.

Wet microcapsules prepared under optimal conditions were lyophilized for 48 h, mounted on conductive tape, and observed by scanning electron microscope (Quanta 450, FEI).

#### Viability of free and encapsulated strain in SGF

One gram of wet microcapsules was weighed and added to 9 mL of SGF, followed by incubation at 37℃ with shaking (220 rpm) for 30, 60, 90, and 120 min. As a control, 1 mL of free (unencapsulated) bacterial suspension was added into 9 mL of SGF, and samples were collected at the same time points under identical conditions. All experiments were performed in triplicate.

#### *In vitro* simulated release assay

One gram of wet microcapsules was weighed and transferred to 9 mL of SIF, then incubated at 37℃ with shaking (220 rpm). Samples were collected at 1, 2, 3, and 4 h. For the control group, 1 mL of free bacterial suspension was added to 9 mL of SIF, and sampling followed the same protocol. Each treatment was repeated three times.

#### Animal and experimental procedure

All experimental procedures and animal care practices in this study adhered strictly to the Guide for the Care and Use of Laboratory Animals established by the Institutional Animal Care and Use Committee of Sichuan Agricultural University. Furthermore, all animal protocols were approved by the same committee under the permit number DKY-B20131704.

A total of 78 healthy 24-day-old piglets with similar body weights were selected and subjected to a 3-day pre-feeding period. One piglet died of illness during pre-feeding, resulting in the following radomization for the formal trial: control group (CON, *n* = 17), T group (basal diet + unencapsulted strain at 1×10^10^ CFU/mL, *n* = 20), TJ group (basal diet + microcapsules of the strain at 1×10^10^ CFU/g, *n* = 20), and J group (basal diet + empty capsules, *n* = 20). The T group received suspension of the strain (saline-based) every other day, while TJ and J groups were supplemented with 1 g of microcapsules or empty capsules, respectively. All groups maintained identical viable strain counts, with supplements administered after feeding. Piglets were fed a corn-soybean meal basal diet free of antibiotics and growth-promoting zinc, meeting the National Research Council (11) nutrient requirements (Table S1). Piglets were fed three times daily (at 8:00 AM, 2:00 PM, and 8:00 PM) with ad libitum access to water. The experiment adhered to animal welfare gauidelines, with controlled environmental conditions (24-26℃, 55-65% humidity). The experiment lasted 40 days.

Individual body weights were recorded the start and end of the trial, with daily feed intake documented to calculate average daily feed intake (ADFI), average daily gain (ADG), and feed-to-gain ratio (F:G). Diarrhea was scored three time daily (8:00, 12:00, and 16:00) using a 0-3 scale: 0 (formed feces), 1 (soft feces), 2 (sticky/liquid), 3 (watery).

The diarrhea index was calculated as:

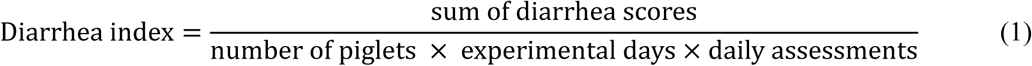

Rectal fecal samples were collected on day 36-39, fixed with 10% sulfuric acid, preserved with toluene, and stored at −20°C. The method of nutrient apparent digestibility based on classical analytical method (12). On day 40, piglets were fasted overnight. Five per group were randomly selected, anesthetized with sodium pentobarbital (200 mg/kg), and euthanized. Intestinal segments were isolated: mid-jejunum fixed in 4% paraformaldehyde for histomorphology, and proximal colon placed in Carnoy’s fixative for mucus/goblet cell analysis. Jejunal and colonic mucosal scrapings were collected, snap-frozen in liquid nitrogen, and stored at −80°C for gene expression analysis. Colonic digesta were collected, flash-frozen in liquid nitrogen, and stored at −80°C for microbial enumeration and SCFAs analysis.

#### Determination of blood biochemical indices

Serum concentrations of cytokines, including interleukin 10 (IL-10) and tumor necrosis factor-α (TNF-α), and immunoglobulins (IgA and IgG) were measured using porcine-specific ELISA kits (Procine ELISA KIT, Quanzhou Ruixin Bio-technology Co., Ltd.). Assays were performed according to the manufacturer’s instructions with strict and adherence to protocol specifications.

#### Intestinal histomorphological analysis

The collected jejunum was fixed in 4% paraformaldehyde solution and processed for paraffin sectioning through dehydration, clearing, and embedding. Sections were cut at 5 μm thickness, stained, mounted, and observed under an inverted microscope (ECLIPSETS100, Nikon, Japan). For each slide, 10 non-overlapping fields of view were randomly selected to measure villus height and crypt depth, from which the villus-crypt ratio was calculated.

Colon samples fixed in Carnoy’s solution (ethanol : chloroform : acetic acid = 6:3:1, v/v) and stored at 4°C were dehydrate with absolute ethanol (two times, 40-60 min each), cleared with xylene, and embedded in paraffin. Paraffin sections were stained with Alcian blue-nuclear fast red (AB-PAS) to visualize the mucus layer, then observed using an Olympus FV1000 laser confocal microscope (Olympus, Japan). Images of colonic goblet cells were captured with a Cap Studio-SC600C camera, and cell counts were performed using ImageJ software.

#### Quantification of intestinal barrier related genes and specific bacterial populations

Total RNA was extracted from ileal mucosa using RNAiso Plus (Takara Bio Inc., Japan), and RNA concentration and purity were assessed with a nanospectrophotometer (NanoDrop2000, Thermo Fisher Scientific, USA). Subsequently, cDNA was synthesized from RNA using a reverse transcription kit (HiScript IV All-in-One Ultra RT SuperMix for qPCR, Vazyme, China). The genes analyzed included zonula occludens-1 (ZO-1), claudin-1 (claudin1), and occludin, with specific primers listed in Table S2. Real-time time PCR was performed on a Quantstudio 5 (Thermo Fisher Scientific, USA) using a reaction mixture containing 5 μL of SYBR Premix Ex Taq™, 0.4 μL each forward and reverse primer, 1 μL of cDNA, and 3.2 μL of RNase-free water. β-actin and GAPDH served as reference genes for calculating relative gene expression levels via the 2^△△Ct^ method.

For bacterial quantification in colonic digesta, samples were thawed on ice, and 0.2g of each sample was used to extract total DNA with the TIANamp Stool DNA Kit (Tiangen Biochemical Technology (Beijing) Co., Ltd., China) according to the manufacturer’s protocol. Quantitative real-time PCR (qPCR) was performed to assess the abundance of *Bacteroides*, *Lactobacillus*, *Bifidobacterium*, and *Escherichia coli* using specific primers (Table S3).

### Statistical analysis

All data were first sorted using Microsoft Excel 2019 and then analyzed with the One-way ANOVA module in SPSS 27.0 after confirming normality. Intergroup comparison between two groups were performed using Student’s t-test, while multiple group comparisons were conducted with the Bonferroni post-hoc test. Data are presented as mean±standard error of the mean (SEM), and statistical significance was set at *P* < 0.05.

## RESULTS

### Identification of *B. thetaiotaomicron* strain

The 16S rDNA sequence of the isolated strain showed 99.79% similarity to *B. thetaiotaomicron* strain JCM 5827 (NCBI GenBank), consistent with intraspecies divergence among different strains. This porcine isolate was therefore designated *B. thetaiotaomicron* LYH5 (hereinafter referred to as LYH5), as shown in the phylogenetic tree (Fig. 1A).

**Fig. 1.**
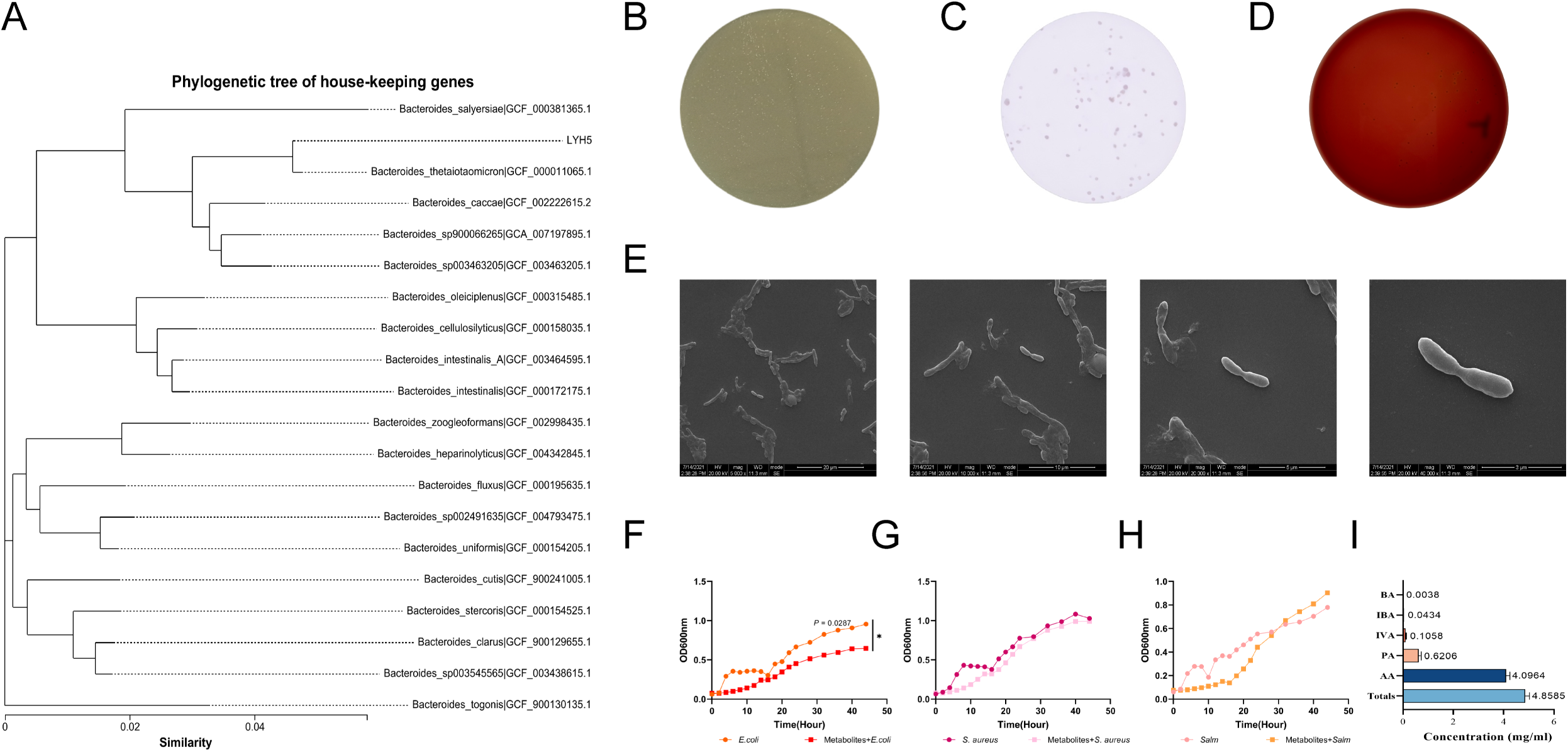
Taxonomic identification and physiological characterization of LYH5. (A) Phylogenetic tree of LYH5 based on 16S rRNA sequence. (B) Colony morphology of LYH5 on modified GAM solid medium. (C) Gram staining of LYH5 (G^-^ rods). (D) Hemolytic activity assay of LYH5 on sheep blood agar (non-hemolytic phenotype). (E) Scanning electron microscopy image of LYH5 cells. (F) Growth curves of *E. coli* ATCC 25922 and *E. coli* ATCC 25922 with LYH5 metabolites (G) Growth curves of *S. aureus* ATCC 25923 and *S. aureus* ATCC 25923 with LYH5 metabolites (H) Growth curves of *S. cholerae suis* ATCC 527 and *S. cholerae suis* ATCC 527 with LYH5 metabolites. (I) SCFAs profile of LYH5 in modified GAM medium. Totals: total SCFAs; AA: acetic acid; PA: propionic acid; IVA: isovaleric acid; IBA: iso-butyric acid; BA: butyric acid.

Morphologically, LYH5 formed round, yellow colonies on solid medium and was identified as a Gram-negative rod (Fig. 1B-C). Physiological characterization revealed catalase-negative, gelatin liquefaction-negative, and non-motile phenotypes (Table S4). Safety evaluations showed non-hemolytic activity (Fig. 1D) and broad antibiotic sensitivity (Table S5).

### *In vitro* bacteriostatic properties of *B. thetaiotaomicron* LYH5 metabolites

The agar diffusion Oxford cup assay (Fig. S1) showed no inhibition zones against *E. coli* ATCC 25922, *S. cholerae suis* ATCC 527, and *S. aureus* ATCC 25923, indicating that viable LYH5 cells do not directly inhibit the growth of these pathogens. However, when co-cultured with extracted LYH5 metabolites (Fig. 1F-H), *E. coli* exhibited significantly reduced growth rates compared to untreated controls (*P* = 0.029).

GC analysis (Fig. 1I) revealed that LYH5 produced approximately 4.8585 mg/mL of total SCFAs in modified GAM medium after 72 h. Acetic acid was the most abundant SCFA, followed by propionic acid, isovaleric acid, isobutyric acid, and butyric acid.

Scanning electron microscopy of LYH5 OMVs isolated via ultracentrifugation confirmed the presence of intact, well-structured vesicles (Fig. S2), indicating that LYH5 actively secretes OMVs under experimental conditions.

### Response surface design and charaterization of *B. thetaiotaomicron* LYH5 microcapsules

Based on single-factor tests results (Fig. S3), Box-Behnken design was used to optimize microencapsulation conditions, with SA (A), gellan gum addition (B), calcium chloride concentration (C), and bacteria-to-gel ratio (D) as independent variables and unit mass encapsulation efficiency as the response variable (Table 2).

**Table 2.** Box-Behnken design matrix and experimental results for optimization of LYH5 microcapsulation parameters.

| Run no. | A | B | C | D | Embedding rate (%) |
| --- | --- | --- | --- | --- | --- |
| 1 | 1 | 0 | -1 | 0 | 65.66 |
| 2 | 0 | 1 | -1 | 0 | 66.78 |
| 3 | 1 | 0 | 0 | -1 | 70.97 |
| 4 | 0 | 1 | 0 | 1 | 71.30 |
| 5 | 0 | 0 | -1 | -1 | 69.22 |
| 6 | 0 | 0 | -1 | 1 | 70.26 |
| 7 | 1 | 0 | 0 | 1 | 70.62 |
| 8 | 0 | -1 | 0 | 1 | 78.09 |
| 9 | -1 | 0 | 1 | 0 | 76.70 |
| 10 | 0 | -1 | 0 | -1 | 77.57 |
| 11 | -1 | -1 | 0 | 0 | 69.57 |
| 12 | 1 | 0 | 1 | 0 | 78.05 |
| 13 | -1 | 0 | -1 | 0 | 67.30 |
| 14 | 0 | 0 | 0 | 0 | 84.70 |
| 15 | 0 | 0 | 0 | 0 | 83.13 |
| 16 | 1 | 1 | 0 | 0 | 69.73 |
| 17 | 0 | 0 | 1 | -1 | 80.70 |
| 18 | -1 | 0 | 0 | 1 | 68.70 |
| 19 | 0 | 0 | 0 | 0 | 82.26 |
| 20 | 0 | -1 | -1 | 0 | 70.09 |
| 21 | -1 | 0 | 0 | -1 | 69.74 |
| 22 | 0 | -1 | 1 | 0 | 82.30 |
| 23 | 1 | -1 | 0 | 0 | 71.50 |
| 24 | 0 | 0 | 1 | 1 | 80.71 |
| 25 | 0 | 0 | 0 | 0 | 83.19 |
| 26 | 0 | 0 | 0 | 0 | 81.24 |
| 27 | 0 | 1 | 0 | -1 | 72.21 |
| 28 | -1 | 1 | 0 | 0 | 63.65 |
| 29 | 0 | 1 | 1 | 0 | 73.98 |

A: SA, B: gellan gum, C: calcium chloride, D: bacteria-to-gel ratio.

Using Design-Expert 10 software, a quadratic regression model was fitted to the data, yielding the equation:

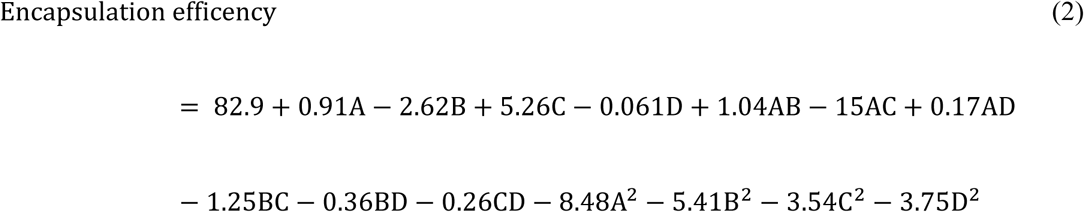

ANOVA for the response surface model is shown in Table 3. Response surface plots (Fig. 2A-F) exhibited downward-opening arches, indicating a maximum encapsulation efficency. The optimal theoretical parameters were determined as: SA 1.481%, gellan gum 0.391%, calcium chloride 0.854%, and bacterial-to-gel ratio 1:4.255, predicting an 84.77% encapsulation efficiency. For pratical application, parameters were adjusted to SA 1.5%, gellan gum 0.4%, calcium chloride 0.9%, and bacterial-to-gel ratio 1:4. Under these conditions, validation experiments showed an average encapsulation efficiency of 84.22±0.17%, highly consistent with the predicted value, confirming the feasibility for response surface methodology for process optimization.

**Fig. 2.**
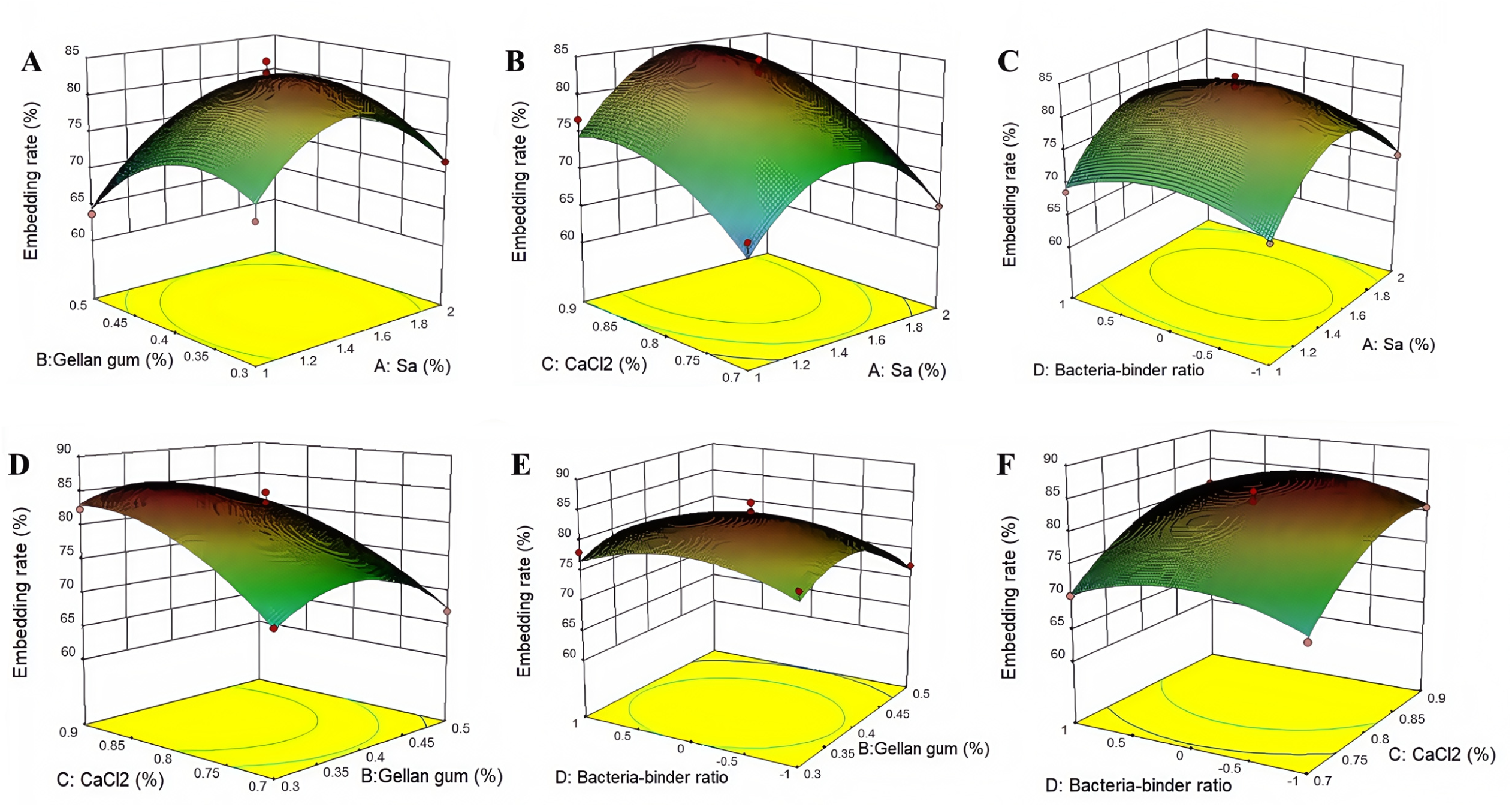
Response surface analysis of process parameters for LYH5 microencapsulation. (A) Relationship between the embedding rate and the concentrations of Gellan gum and Sa. (B) Relationship between the embedding rate and the concentrations of CaCl₂ and Sa. (C) relationship between the embedding rate and the concentrations of the bacteria-binder ratio and Sa. (D) relationship between the embedding rate and the concentrations of CaCl_2_ and Gellan gum. (E) Relationship between the embedding rate and the concentrations of Gellan gum and the bacteria-binder ratio. (F) Relationship between the embedding rate and the concentrations of CaCl_2_ and the bacteria-binder ratio. The 3D surface plots and corresponding contour plots providing a comprehensive visualization of the relationships and interactions among these variables.

**Table 3.** Analysis of variance for response surface model of LYH5 microcapsule encapsulation effiencey.

| Source | Sum of Squares | df | Mean Square | F Value | P-value |
| --- | --- | --- | --- | --- | --- |
| Model | 1027.65 | 14 | 73.40 | 30.65 | < 0.001 |
| A- SA (%) | 9.89 | 1 | 9.89 | 4.13 | 0.062 |
| B- Gellan glue (%) | 82.39 | 1 | 82.39 | 34.40 | < 0.001 |
| C- CaCl2 (%) | 331.99 | 1 | 331.99 | 138.62 | < 0.001 |
| D- Bacteria-gum ratio | 0.04 | 1 | 0.04 | 0.02 | 0.894 |
| AB | 4.29 | 1 | 4.29 | 1.79 | 0.202 |
| AC | 2.25 | 1 | 2.25 | 0.94 | 0.349 |
| AD | 0.12 | 1 | 0.12 | 0.05 | 0.827 |
| BC | 6.28 | 1 | 6.28 | 2.62 | 0.128 |
| BD | 0.51 | 1 | 0.51 | 0.21 | 0.6518 |
| CD | 0.27 | 1 | 0.27 | 0.11 | 0.7448 |
| A^2 | 466.96 | 1 | 466.96 | 194.97 | < 0.001 |
| B^2 | 190.01 | 1 | 190.01 | 79.33 | < 0.001 |
| C^2 | 81.32 | 1 | 81.32 | 33.95 | < 0.001 |
| D^2 | 91.21 | 1 | 91.21 | 38.08 | < 0.001 |
| Residual | 33.53 | 14 | 2.40 |  |  |
| Lack of Fit | 27.00 | 10 | 2.70 | 1.65 | 0.332 |
| Pure Error | 6.53 | 4 | 1.63 |  |  |
| Cor Total | 1061.18 | 28 |  |  |  |

### Tolerance and release kinetics of *B. thetaiotaomicron* LYH5 and its microcapsules in simulated gastrointestinal fluids

Uncapsulated LYH5 showed rapid viability decline in SGF, with survival rates dropping to 12% after 1 h (Fig. 3A). In contrast, viability remained stable in SIF, maintaining 71% survival after 1 h. Microencapsulated LYH5 exhibited significantly enhanced SGF tolerance (Fig. 3B). Uncapsulated LYH5 showed survival rates of 18% at 60 min and 6% at 120 min, whereas microencapsulated LYH5 retained 65% viability after 120 min. Enteric-coated microcapsules demonstrated controlled bacterial release in SIF (Fig. 3C). Viable cell counts increased from 2.94 lg(CFU/g) at 1 h to 3.61 lg(CFU/g) at 4 h, indicating sustained release in the intestinal environment.

**Fig. 3.**
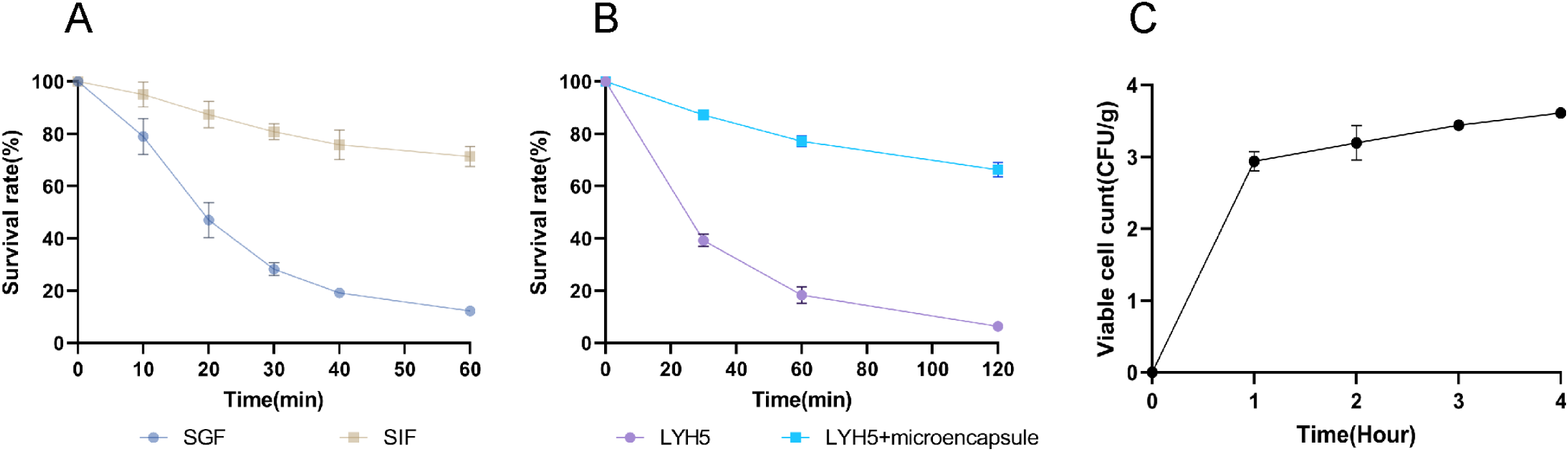
Tolerance and release kinetics of LYH5 microcapsules in simulated gastrointestinal fluids. (A) Viability assay of LYH5 in SGF. (B) Comparative survival rates of uncapsulated and microencapsulated LYH5 in SGF. (C) Time-dependent release kinetics of LYH5 microcapsules in SIF.

### Impact of *B. thetaiotaomicron* LYH5 microcapsules on phenotypic parameters of weaned piglets

During the entire experimental period, dietary supplementation with LYH5 microcapsules did not affect ADG, ADFI, F:G, or diarrhea index in weaned piglets compared to the CON group (*P* > 0.05, Table 4). The TJ group exhibited a trend toward lower F:G relative to the CON group (*P* = 0.071).

**Table 4.**
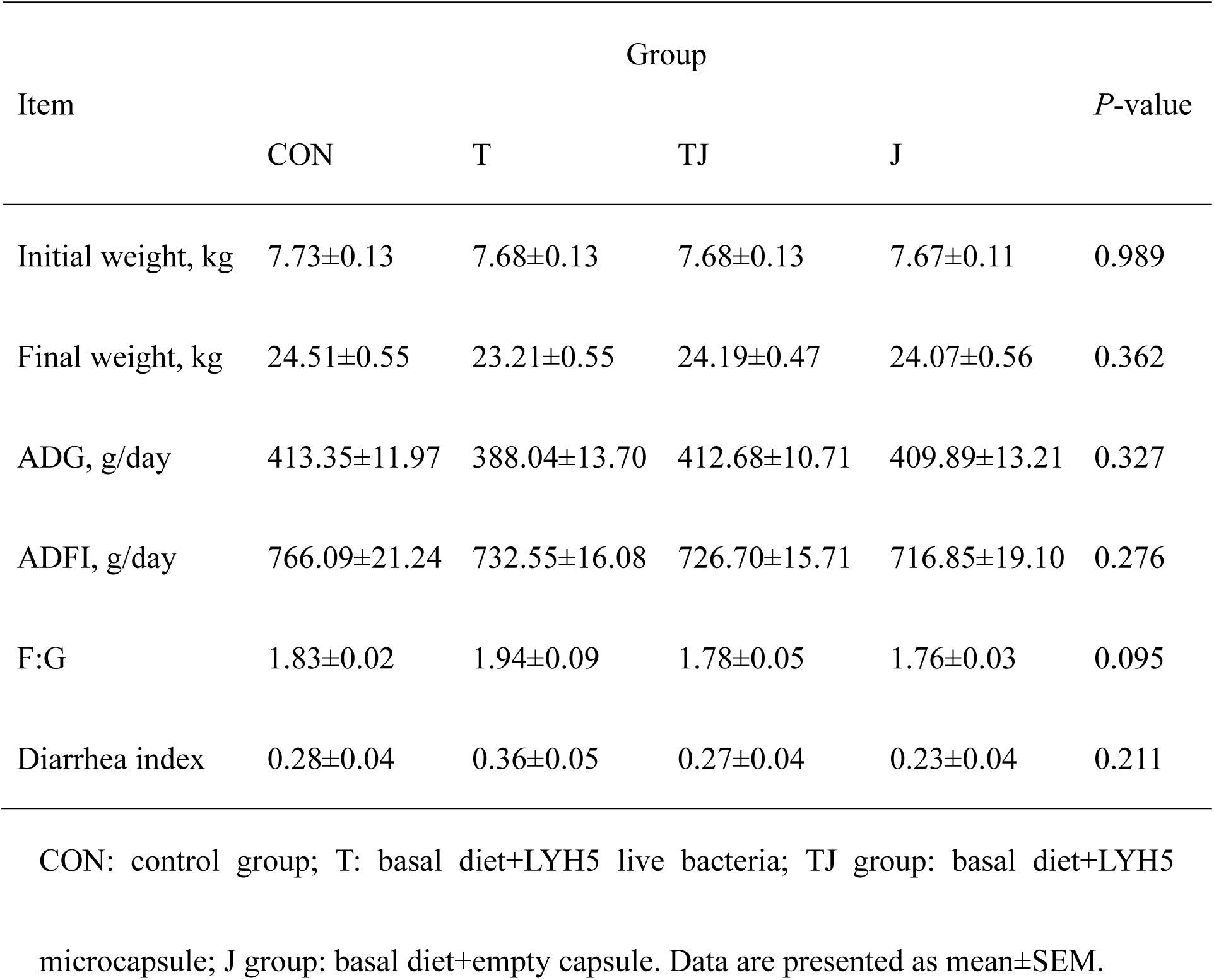
Comparison of growth performance and diarrhea incidence in weaned piglets in different groups.

| Item | Group |  |  |  | *P*-value |
| --- | --- | --- | --- | --- | --- |
|  | CON | T | TJ | J |  |
| Initial weight, kg | 7.73±0.13 | 7.68±0.13 | 7.68±0.13 | 7.67±0.11 | 0.989 |
| Final weight, kg | 24.51±0.55 | 23.21±0.55 | 24.19±0.47 | 24.07±0.56 | 0.362 |
| ADG, g/day | 413.35±11.97 | 388.04±13.70 | 412.68±10.71 | 409.89±13.21 | 0.327 |
| ADFI, g/day | 766.09±21.24 | 732.55±16.08 | 726.70±15.71 | 716.85±19.10 | 0.276 |
| F:G | 1.83±0.02 | 1.94±0.09 | 1.78±0.05 | 1.76±0.03 | 0.095 |
| Diarrhea index | 0.28±0.04 | 0.36±0.05 | 0.27±0.04 | 0.23±0.04 | 0.211 |

CON: control group; T: basal diet+LYH5 live bacteria; TJ group: basal diet+LYH5 microcapsule; J group: basal diet+empty capsule. Data are presented as mean±SEM.

For apparent nutrient digestibility, LYH5 microcapsules did not impact nutrient digestibilities in weaned piglets (*P* > 0.05, Table S6), though the T group showed a tendency for increased crude protein digestibility (*P* = 0.093).

Regarding blood biochemical indices (Table S7), serum free fatty acid levels in the TJ group trended higher than the CON group (*P* = 0.083), but no significant differences were observed on other blood parameters across treatment groups (*P* > 0.05).

### Impact of *B. thetaiotaomicron* LYH5 microcapsules on serum cytokines and immunoglobulin

Serum IgG concentration in the CON group was significantly lower than that in the TJ and J groups (*P* < 0.05, Fig. 4A), with the TJ group exhibiting the highest level at 1.73 g/L. The T group showed a trend toward increased serum IL-10 concentration compared to CON (*P* = 0.092, Fig. 4D). No differences were observed in serum IgA and TNF-α levels across groups (*P* > 0.05, Fig. 4B-C).

**Fig. 4.**
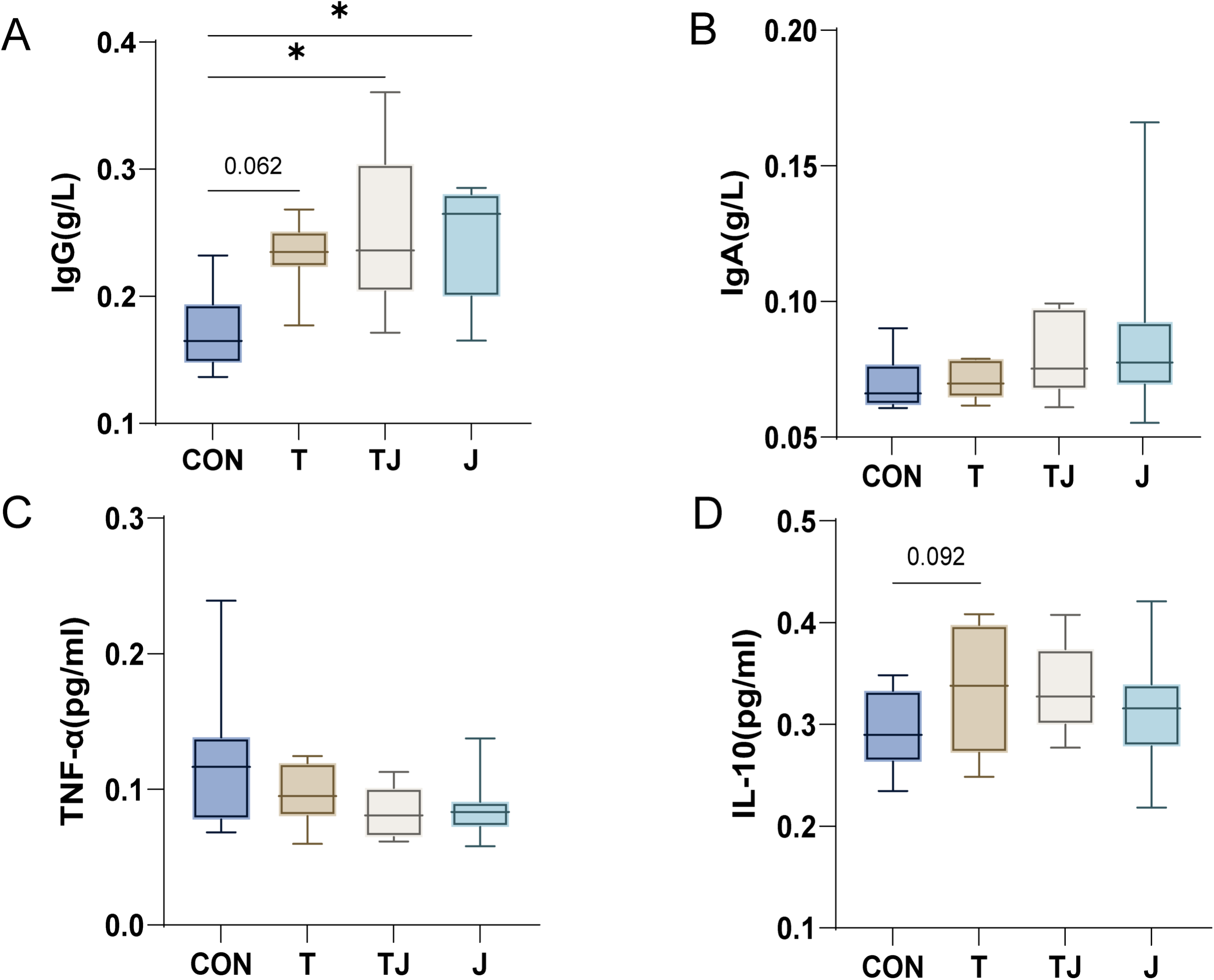
LYH5 microcapsules modulate serum cytokines and immunoglobulins in weaned piglets. (A) Serum IgG. (B) Serum IgA. (C) Serum TNF-α. (D) Serum IL-10. Data are presented as mean±SEM. Statistical significance: \**P* < 0.05.

### Effects of *B. thetaiotaomicron* LYH5 microcapsules on jejunal morphology and colonic mucus layer in weaned piglets

Compared to the CON group, the TJ group exhibited greater jejunal villus height (*P* < 0.001) and villus height-to-crypt depth ratio (*P* < 0.001, Fig. 5A-D). No significant differences were noted between the J and CON groups (*P* > 0.05). The TJ group also showed a thicker colonic mucus layer than both the CON group (*P* < 0.001) and T group (*P* = 0.024, Fig. 5E), along with a trend toward increased colonic goblet cell counts (*P* = 0.048, Fig. 5F).

**Fig. 5.**
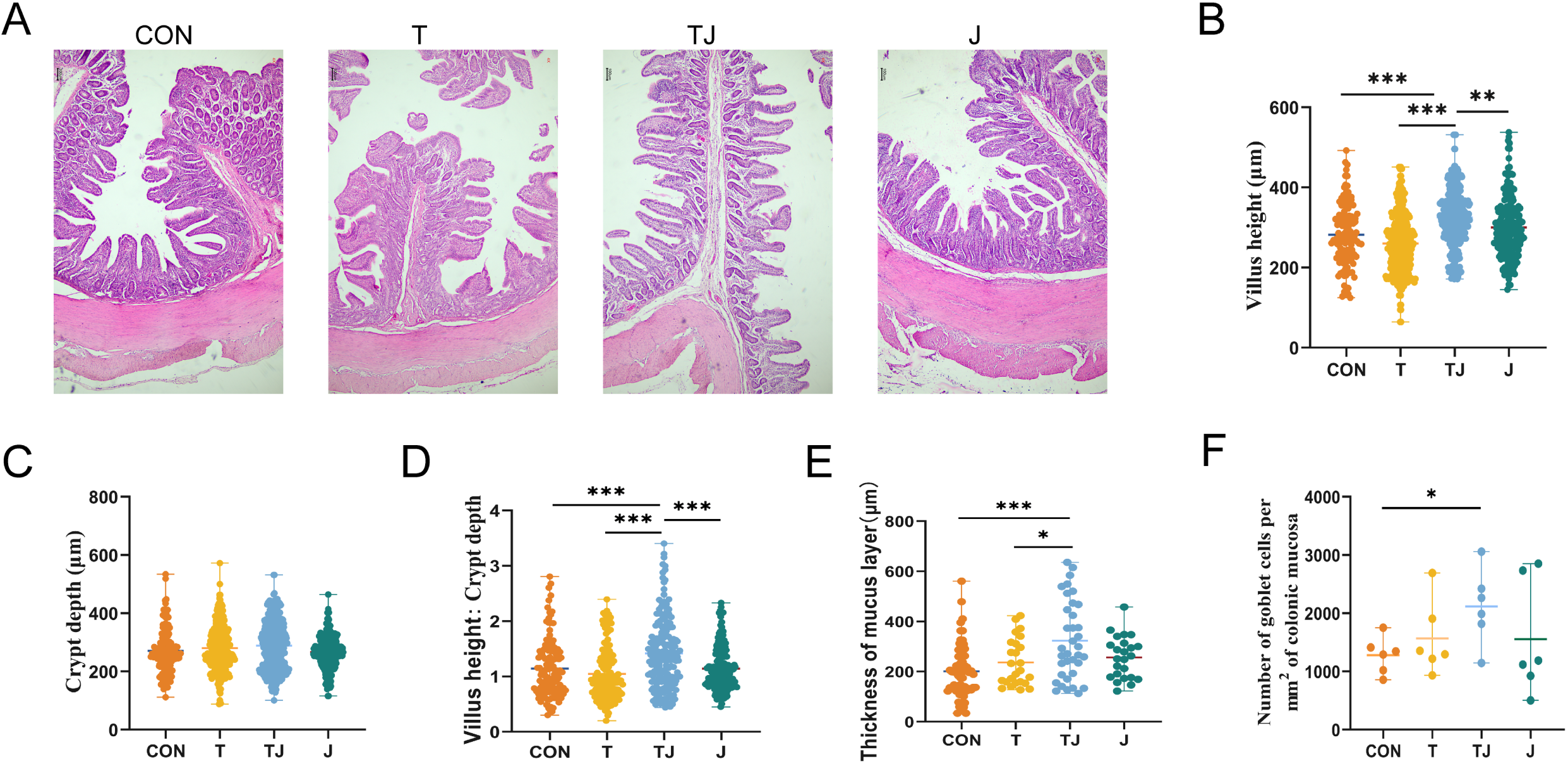
LYH5 microcapsules improve jejunal morphology and colon mucus layer thickness in weaned piglets. (A) Jejunal histomorphology (HE staining). (B) Villus height. (C) Crypt depth. (D) Villus height-to-crypt depth ratio. (E) Colonic mucus layer thickness. (F) Colonic goblet cell cout. Data are shown as mea ±SEM. Statistical significance: \**P* < 0.05, \*\**P* < 0.01, \*\*\**P* < 0.001.

### Effects of *B. thetaiotaomicron* LYH5 microcapsules on intestinal barrier related gene expression and certain bacterial groups in colonic digesta of weaned piglets

In the jejunum, the T group exhibited upregulated ZO-1 expression compared to the CON group (*P* = 0.005, Fig. 6A), along with trends toward increased Claudin-1 (*P* = 0.012, Fig. 6C) and Occludin (*P* = 0.071, Fig. 6D), while jejunal ZO-2 expression remained unchanged across all groups (*P* > 0.05, Fig. 6B). In the colon, the TJ group showed higher ZO-2 (*P* = 0.007 vs. CON, *P* = 0.003 vs. T, Fig. 6F) and Claudin-1 (*P* = 0.012 vs. CON, *P* = 0.003 vs. T, Fig. 6G) expression than both the CON and T groups.

**Fig. 6.**
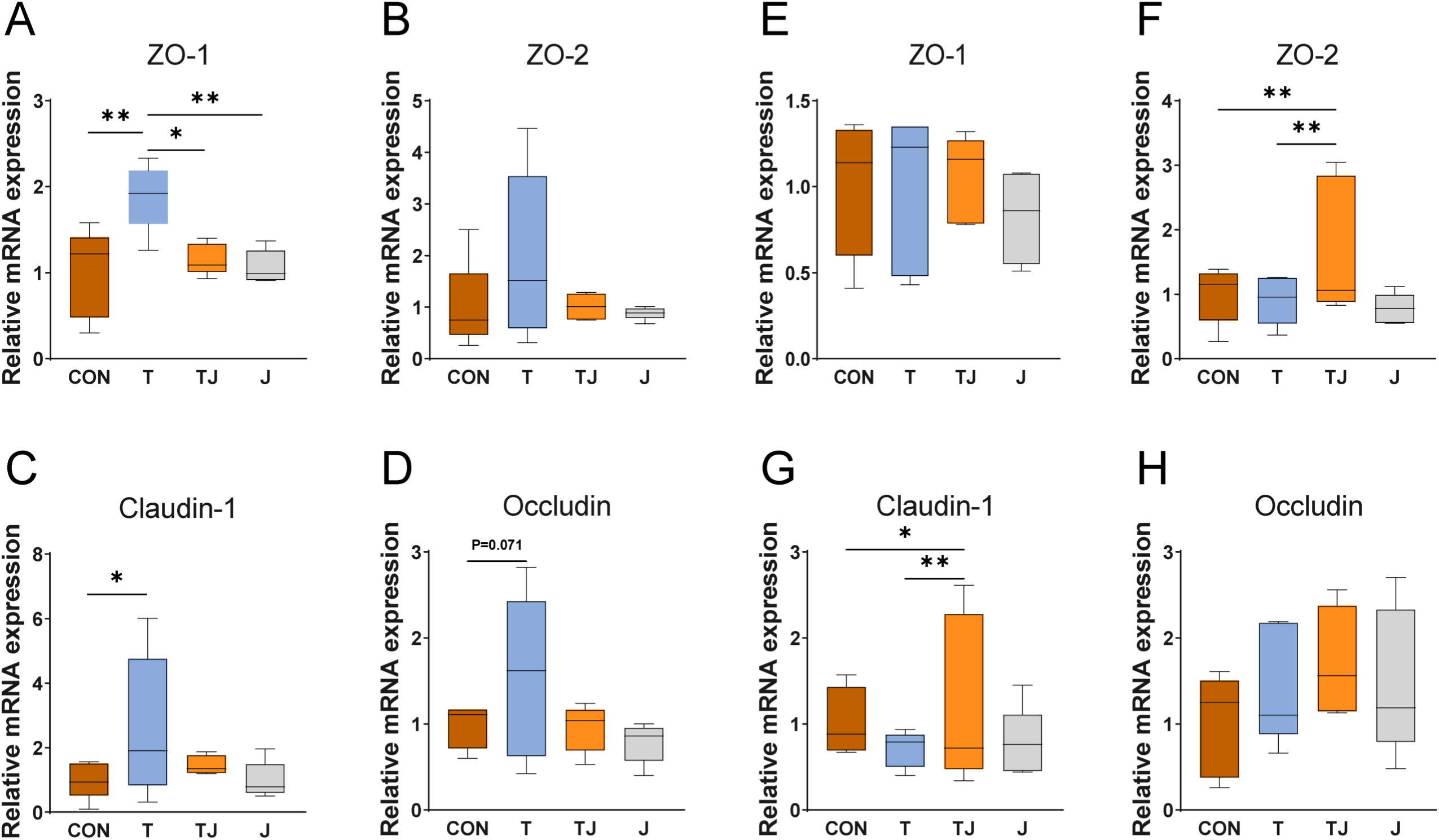
LYH5 microcapsules modulate tight junction gene expression in jejunum and colon of weaned piglets. (A) Relative mRNA expression of ZO-1 in the jejunum. (B) Relative mRNA expression of ZO-2 in the jejunum. (C) Relative mRNA expression of Claudin-1 in the jejunum. (D) Relative mRNA expression of Occludin in the jejunum. (E) Relative mRNA expression of ZO-1 in the colon. (F) Relative mRNA expression of ZO-2 in the colon. (G) Relative mRNA expression of Claudin-1 in the colon. (H) Relative mRNA expression of Occludin in the colon. Data are presented as mean±SEM. Statistical significance: \**P* < 0.05, \*\**P* < 0.01.

Results of qPCR targeting specific bacterial groups (Table 5) showed total bacterial counts in the J group were lower than in the CON (*P* = 0.048), T (*P* = 0.010) and TJ (*P* = 0.039) groups. *Bacteroides* and *Lactobacillus* numbers were similar across all groups (*P* > 0.05), while the TJ group had higher *Bifidobacterium* counts than both the T (*P* = 0.005) and CON (*P* = 0.029) groups. *E. coli* numbers in the T (*P* = 0.016) and TJ (*P* = 0.029) groups waslower than that in the CON group.

**Table 5.**
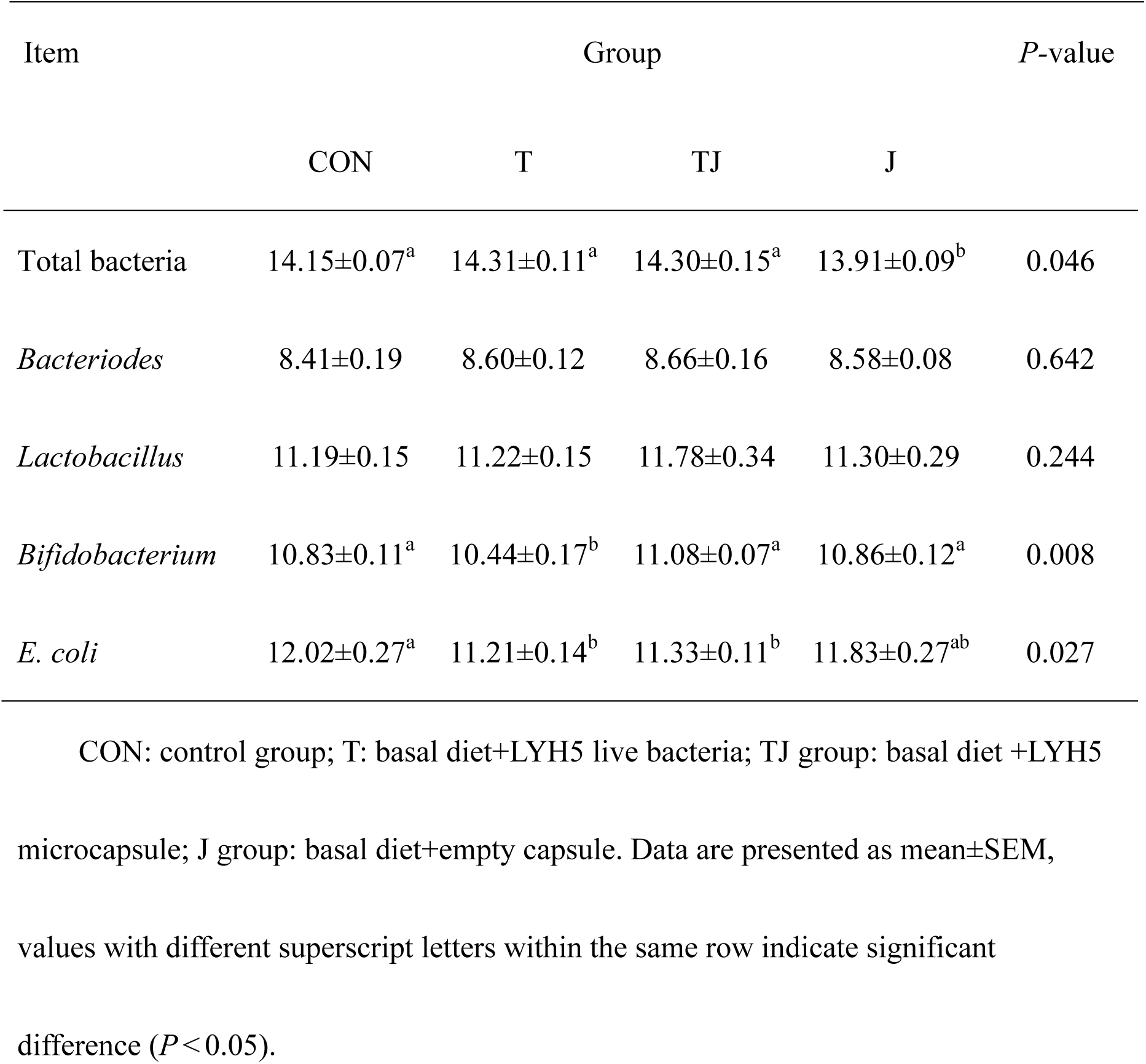
Comparison of certain bacterial group numbers in colonic digesta of weaned piglets in different groups [lg (copies)/g wet digesta].

|  | CON | T | TJ | J |  |
| --- | --- | --- | --- | --- | --- |
| Total bacteria | 14.15±0.07<sup>a</sup> | 14.31±0.11<sup>a</sup> | 14.30±0.15<sup>a</sup> | 13.91±0.09<sup>b</sup> | 0.046 |
| *Bacteriodes* | 8.41±0.19 | 8.60±0.12 | 8.66±0.16 | 8.58±0.08 | 0.642 |
| *Lactobacillus* | 11.19±0.15 | 11.22±0.15 | 11.78±0.34 | 11.30±0.29 | 0.244 |
| *Bifidobacterium* | 10.83±0.11<sup>a</sup> | 10.44±0.17<sup>b</sup> | 11.08±0.07<sup>a</sup> | 10.86±0.12<sup>a</sup> | 0.008 |
| *E. coli* | 12.02±0.27<sup>a</sup> | 11.21±0.14<sup>b</sup> | 11.33±0.11<sup>b</sup> | 11.83±0.27<sup>ab</sup> | 0.027 |

CON: control group; T: basal diet+LYH5 live bacteria; TJ group: basal diet +LYH5
microcapsule; J group: basal diet+empty capsule. Data are presented as mean±SEM,
values with different superscript letters within the same row indicate significant
difference ( $P < 0.05$ ).

## DISCUSSION

Despite the growing interest in *Bacteroides* species as probiotics, previous studies have primarily focused on their immunomodulatory and gut microbiota-modulating effects, with limited investigation into their biosafety and functional metabolites for practical applications in livestock nutrition. In this study, LYH5 was identified as a novel *B. thetaiotaomicron* strain based on 16S rRNA sequence analysis, morphological characteristics, and physiological-biochemical properties. The absence of hemolytic activity and sensitivity to most antibiotics further support its potential as a safe probiotic candidate. These findings align with previous investigations on *Bacteroides* species, which have been widely recognized for their probiotic potential in improving gut health and modulating immune responses(5, 14). The high biosafety and functional attributes of LYH5 render it a promising candidate for oral supplementation in livestock.

In nature, all organisms rely on their innate immune defense system, encompassing pattern recognition receptors, cytokines, complement cascades, and host defense peptides (HDPs) (41). HDPs, often termed “natural antimicrobial peptides”, exhibit broad-spectrum antibacterial activity against various pathogens (36, 47). Dietary components such as SCFAs, probiotics, and prebiotics can enhance endogenous HDP synthesis without inducing inflammation or compromising barrier integrity (52). The antimicrobial activity of LYH5 was evaluated *in vitro*. Ethyl acetate-extracted metabolites from LYH5 exhibited pronounced effects against *E. coli*, suggesting that its antimicrobial activity is mediated by metabolites (e.g. acetic acid), which may disrupt pathogen cell membranes (40) or interfere with metabolic pathways (39). These findings imply that LYH5 may produce HDPs with biological activity, warranting further mechanistic investigation.

Tolerance to gastric and intestinal fluid is a critical criterion for probiotic evaluation. In this study, the survival rate of LYH5 decreased significantly after SGF treatment, indicating poor resistance to the low pH and digestive enzymes of the stomach. By contrast, its survival rate remained stable SIF treatment, demonstrating its capacity for intestinal adaptability. Given its intolerance to SGF, practical application of LYH5 may require protective strategies, such as appropriate embedding techniques, to ensure its released in the intestine and subsequent physiological effects. Screening and optimizing embedding conditions for this strain thus paves the way for its translational application in livestock production.

Embedding technology has been widely used in food and feed industry. Previous research indicates that a 1% SA concentration offers suboptimal protection for probiotics due to inadequate tolerance (8). Moreover, SA-based microcapsules typically exhibit porous surface, rendering them vulnerable to the low-pH gastric environment. To mitigate this limitation, SA is often combined with other wall materials (4). For instance, *Lactobacillus helveticus* and *Lactobacillus delbrueckii* microcapsules formulated with 2% SA and 0.2% gellan gum mixture have demonstrated sufficient bacterial activity to meet the industry-standard dosage range of 10^6^-10^7^ CFU/g (46). Additionally, spherical microcapsules have been shown to outperform non-spherical counterparts in maintaining integrity within SGF, as non-spherical structures tend to rupture prematurely and release their contents (33). Notably, the LYH5 microcapsules prepared in this study, utilizing SA and gellan gum as wall materials, exhibited a pore-free, uniform, and spherical morphology, suggesting enhanced stability.

Despite the emergence of advanced microcapsule fabrication techniques, the traditional extrusion method employed herein offers distinct advantages,including technical simplicity, cost-effectiveness, and high encapsulation efficiency. Given LYH5’s inherent susceptibility to SGF, microencapsulation serves as a crucial strategy to preserve bacterial activity during gastrointestinal transit. Studies have demonstrated that microcapsules prepared via extrusion, such as that *Salmonella* phage microcapsules, can achieve high microencapsulation efficiency, withstand low-pH conditions, and facilitate targeted release in SIF (19). Correspondingly, the LYH5 microcapsules achieved an encapsulation rate of approximately 84.22%, exhibited notable tolerance to SGF, and enabled effective released in SIF. These findings underscore the practical utility of the developed microcapsules, providing a viable solution for translating LYH5 into livestock probiotic application.

While previous research has predominantly explored the impact of *Bacteroides* on physiological phenotypes within obesity models, findings have been equivocal. Epidemiological studies in both adult and pediatric populations have consistently reported reduced gut *Bacteroides* abundance in obese individuals (31, 48). Conversely, recent investigations have revealed a positive correlation between *B. thetaiotaomicron* levels and body mass index across both metabolically healthy and unhealthy obesity phenotypes (2), challenging the simplistic notion of a negative association between *Bacteroides* and obesity. Mechanistic studies further underscore the metabolic benefits of specific *Bacteroides* strains. For example, *B. uniformis* 7771 administration mitigated metabolic dysfunction in obese mice by reducing serum triglycerides, cholesterol, and glucose levels (20), while *Bacteroides* supplementation reversed weight loss associated with intestinal disease (15). Notebly, our study did not detect significant effects of LYH5 microcapsules on the growth performance or apparent nutrient digestibility of healthy weaned piglets. This discrepancy may be attributed to the contrasting experimental contexts. Prior research predominantly utilized disease models (e.g. obesity or enteritis), highlighting traom-specific anti-inflammatory properties, whereas our study focused on healthy subjects. Nevertheless, the absence of adverse effects on piglet growth validates a crucial prerequisite for LYH5’s potential use as a probiotic.

Immunoglobulins, a class of structurally globulins secreted by lymphocytes, mediate diverse immune functions, including antimaicrobial activity, antiviral defense, immune modulation, and complement activation (24, 55). Among these, IgG represents the most abundant immunoglobulin in animal serum, constituting 75%-80% of total antibodies (9). Previous investigations have demonstrated that dietary probiotic supplementation can elevate serum IgG concentrations in piglets (32), potentially reducing intestinal pathogen colonization and preventing enteric diseases (26). Extracellular vesicles (EVs) derived from human *B. thetaiotaomicron* have been shown to maintain immune homeostasis by interacting with innate immune cells and eliciting anti-inflammatory responses (22). Intriguingly, our study identified OMV production by the porcine-derived *B. thetaiotaomicron* strain, suggesting a potential role in modulating host immune function. However, further validation in pathological models is essential to elucidate the immunomodulatory mechanisms of LYH5-derived EVs.

IL-10 is a pivotal cytokine in the regulation of inflammation and immune responses, serving as a key modulator of host inflammatory reactions during infections with diverse pathogens (23). It exerts its immunomodulatory effects by inhibiting the synthesis of pro-inflammatory factors and chemotaxis while simultaneously promoting the secretion of anti-inflammatory cytokines (18). Previous investigations have established associations between increased gut *Bacteroidetes* abundance and elevated serum levels of immunoglobulins and cytokines, alongside reduced TNF-α concentrations in piglets (7, 58). For instance, oral administration of *B. vulgatus* FTJS7K1 upregulated IL-10 expression in LPS-induced mice (51). In line with these findings, our study demonstrated that dietary supplementation with porcine-derived *B. thetaiotaomicron* LYH5 microcapsules or live bacterial solutions significantly increased serum IgG levels in weaned piglets. Moreover, the live bacterial solution exhibited a trend towards enhancing IL-10 production, suggesting that LYH5 may promote immune function in piglets. However, additional research using pathological models is warranted to elucidate the precise mechanisms underlying LYH5-mediated immune enhancement.

The integrity of intestinal morphology is fundamental to maintaining normal digestive and absorptive functions, with villus height and crypt depth serving as critical biomarkers for assessing intestinal health. As extensively documented, a higher villus height-to-crypt depth ratio typically correlates with enhanced nutrient digestion and absorption capabilities (27). Given the incomplete intestinal development in weaned piglets, interventions that improve intestinal morphology can promote their physiological functions. Despite the growing interest in NGPs, limited research has explored their impact on intestinal morphology. Existing studies have shown strain-specific effects. In a neonatal rat model of necrotizing enterocolitis induced by *Cronobacter sakazakii*, treatment with *B. fragilis* ZY-312 mitigated intestinal injury by preserving villus height (21), while other *B. fragilis* strains, such as FSHCM14E1 and NCTC9343, restored colon morphology in mice with DSS-induced colitis (50). Conversely, *B. fragilis* JSWX11BF failed to alleviated DSS-induced colitis (21), highlighting the variability in probiotic efficacy. Notably, our study revealed that feeding LYH5 microcapsules, but not unencapsulated live bacteria or emty capsules, significantly increased jejunal villus height and improved the villus height-to-crypt depth ratio in healthy weaned piglets. These results suggest that microencapsulation is essential for LYH5 to exert its beneficial effects on intestinal morphology, providing novel insights into the application of *Bacteroides* as probiotics in swine nutrition.

The intestinal chemical barrier, a pivotal component of the gut mucosal system, is primarily composed of the mucus layer, gastric acid, lysozyme, bile salts, mucoproteins, and antimicrobial peptides (45). These components synergistically exert bactericidal effects and safeguard against the invasion of toxic substances. Goblet cells, specialized epithelial cells lining the intestinal and other mucosal surfaces, are integral to maintaining this barrier by secreting mucus, which forms a protective gel layer overlying the epithelium (13). Previou studies have underscored the role *Bacteroides* strains in modulating goblet cell function and mucus production. For instance, oral administration of *B. vulgatus* 7K1 increases the number of goblet cells in the colons of DSS-treated mice (35). Similarly, *B. fragilis* ZY-312 enhances mucus secretion and intestinal barrier integrity in mice via the signal transducer and activator of transcription (STAT3) signaling pathway (59). Moreover, treatment with *B. ovatus* ATCC 8483 promotes the proliferation of intestinal epithelial cells, goblet cell differentiation, and mucin production in DSS-induced mice, outperforming traditional fecal transplantation in efficacy (29).

In the current study, dietary supplementation with LYH5 microcapsules increased the thickness of the colonic mucus layer in healthy weaned piglets and demonstrated a trend towards augmenting the number of colonic goblet cells. Compared with unencapsulated live bacterial solutions, microcapsules exerted more pronounced effects, indicating that the LYH5 strain has the potential to fortify the intestinal chemical barrier of weaned piglets.

Complementing the chemical barrier, the intestinal physical barrier is fundamental to gut homeostasis. Composed of intact intestinal epithelial cells and intercellular tight junctions, it serves as the structural basis for maintaining the selective permeability of the intestinal epithelium and preserving barrier integrity (49). Tight junctions are complex molecular assemblies comprising transmembrane proteins (occludin, claudins) and intracellular adaptor proteins (ZO-1, ZO-2, ZO-3) that regulate paracellular transport (42). Although research on *Bacteroides*-mediated physical barrier maintenace remains limited, emerging evidence suggests strain-specific benefits. For example, *B. fragilis* ZY-312 reduces the morbidity and mortality of *Clostridium difficile*-infected mice and restores the expression levels of ZO-1 and MUC-2 in the intestinal epithelium (16). The same strain also protects against *Cronobacter sakazakii* infection by upregulating ZO-1 expression and mitigating intestinal damage (21). Our findings revealed differential effects of LYH5 on the physical barrier: the live bacteria more effectively enhanced the gene expression of jejunal tight-junction proteins in healthy weaned piglets, whereas only LYH5 microcapsules promote tight-junction protein gene expression in the colon. These discrepancies likely stem from the unique properties of the microencapsulation process.

*B.thetaiotaomicron* predominantly colonizes the posterior intestine of pigs, where it exerts beneficial effects on gut homeostasis. Given that LYH5 microcapsules exhibit slow-release characteristics in intestinal fluid, their distribution activity may vary along the gastrointestinal tract, potentially resulting in reduced release in the jejunum and enhanced release in the colon. Intriguingly, quantification of colonic bacteria did not reveal an expected increase in *Bacteroides* abundance. This discrepancy may be primarily attributed to the dynamic equilibrium of the gut microbiota in healthy animals, where potential elution mechanisms likely counteract the accumulation of exogenous probiotic strains. To precisely elucidate the colonization patterns of LYH5 within the piglet intestine, future investigations should leverage advanced techniques such as metagenomic sequencing or fluorescence in situ hybridization, which offer higher taxonomic resolution and spatial localization capabilities. Notably, the colonic digesta of the treatment groups (TJ and T) exhibited a significant reduction in E. coli counts compared to the control group. This finding provides additional evidence supporting the probiotic properties LYH5, as the strain likely exerts antimicrobial effects through the production of metabolites or modulation of the gut microenvironment. These results align with previous in vitro observations of LYH5’s inhibitory activity against *E. coli* and underscore its potential to improve gut health by suppressing pathogenic bacteria in the porcine intestine.

## CONCLUSION

In conclusion, we successfully ehnhanced the tolerance of porcine-derived *B. thetaiotaomicron* LYH5 strain to SGF through microencapsulation, ensuring its viability post-gastric transit. In a pig model, dietary supplementation with LYH5 microcapsules significantly elevated serum of IgG levels, upregulated the gene expression of jejunal tight junction protein ZO-1 and cladin-1 in the colon. These effects occurred without compromising piglet growth performance or nutrent digestibility. Additionally, LYH5 mirocapsules promoted the proliferation of colonic goblet cells and thickened the mucus layer, thereby strengthening the intestinal barrier function. Overall, this research presents an effective strategy for improving the gastric stability of probiotics and offers valuble insights into the development of NGPs targeting intestinal health in young animals. The findings lay a solid foundation for translating *B. thetaiotaomicron*-based probiotics into practical applications in veterinary medicine and livestock agriculture.

## CRediT AUTHORSHIP CONTRIBUTION STATEMENT

**Shen Jin**: Writing - original draft, Visualization, Software, Methodology, Investigation, Formal analysis, Data curation, Conceptualization. **Yanjiao Liu**: Writing - review & editing, Data curation. **Yifang Zhang**: Writing - review & editing, Software. **Yuqing Shen**: Methodology, Investigation. **Cong Lan**: Visualization, Software. **Hua Li**: Visualization, Methodology, Investigation, Funding acquisition. **Jun He**: Visualization, Validation. **Aimin Wu**: Validation, Supervision. **Jiayong Tang**: Methodology, Investigation. **Ruinan Zhang**: Supervision, Resources. **Huifen Wang**: Project administration, Methodology. **Quyuan Wang**: Visualization. **Gang Tian**: Methodology, Investigation. **Jingyi Cai**: Software, Methodology. **Xiangbing Mao**: Project administration, Investigation. **Edward L Good**: Writing - review & editing, Visualization. **Yuheng Luo**: Writing - review & editing, Project administration, Methodology, Investigation, Funding acquisition.

## ACKNOWLEDGMENTS

We thank our colleagues for their help in collecting the data. This study was funded by National Key Research and Development Program (2023YFD1301400 and 2023YFD1301402), and National Natural Science Foundation of China (32372900).

## DATA AVAILABILITY

Correspondent authors upon reasonable requests will provide raw data supporting the findings of this study.

